# A divisive model of evidence accumulation explains uneven weighting of evidence over time

**DOI:** 10.1101/598789

**Authors:** Waitsang Keung, Todd A. Hagen, Robert C. Wilson

## Abstract

Divisive normalization has long been used to account for computations in various neural processes and behaviours. The model proposes that inputs into a neural system are divisively normalized by the total activity of the system. More recently, dynamical versions of divisive normalization have been shown to account for how neural activity evolves over time in value-based decision making. Despite its ubiquity, divisive normalization has not been studied in decisions that require evidence to be integrated over time. Such decisions are important when we do not have all the information available at once. A key feature of such decisions is how evidence is weighted over time, known as the integration ‘kernel’. Here we provide a formal expression for the integration kernel in divisive normalization, and show that divisive normalization can quantitatively account for the perceptual decision making behaviour of 133 human participants, performing as well as the state-of-the-art Drift Diffusion Model, the predominant model for perceptual evidence accumulation.

## Introduction

Divisive normalization has been proposed as a canonical computation in the brain [1]. In these models, the firing rate of an individual neuron is computed as a ratio between its response to an input and the summed activity of a pool of neurons receiving similar inputs. For example, activity of a visual cortex neuron *f*_*i*_ responding to an input *u*_*i*_ can be computed as the input divided by a constant *S* plus a normalization factor — the sum of inputs received by the total pool of neurons [1]:

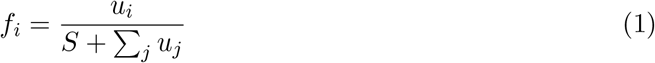

Divisive normalization models such as described in equation 1 have been used successfully to describe both neural firing and behavior across a wide range of tasks — from sensory processing in visual and olfactory systems [2–5], to context-dependent value encoding in premotor and parietal areas [6]. For example, in the visual domain, divisive normalization explains surround suppression in primary visual cortex, where the response of a neuron to a stimulus in the receptive field is suppressed when there are additional stimuli in the surrounding region [7]. Analogously, in economic decision making, divisive normalization explains how activity in parietal cortex encodes the value of a choice option relative to other available alternatives instead of the absolute value [6]. More recently, *dynamic* divisive normalization models have been used to describe how neural activity in economic decision making tasks evolves over time [8, 9].

Despite the success of divisive normalization models, they have never been studied in situations that require evidence to be *integrated* over time. Such ‘evidence accumulation’ is important in many decisions when we do not have all the information available at once, such as when we integrate visual information from moment to moment as our eyes scan a scene.

In the lab setting, evidence accumulation has typically been studied in perceptual decision making tasks over short periods of time. In one such task, called the Poisson Clicks Task [10], participants make a judgment about a train of auditory clicks. Each click comes into either the left or right ear, and at the end of the train of clicks participants must decide which ear received more clicks. The optimal strategy in this task is to ‘count,’ i.e. integrate, the clicks on each side and choose the side with the most clicks.

A key feature of any evidence accumulation strategy is how evidence is weighted over time, which is also known as the ‘kernel’ of integration. For example in the optimal model of counting, each click contributes equally to the decision, i.e., all clicks are weighed equally over time. In this case, the integration kernel is flat — the weight of every click is the same. While such flat integration kernels have been observed in rats and highly trained humans [10], there is considerable variability across species and individuals. For example, [11] showed that monkeys exhibit a strong primacy kernel, in which early evidence is over weighed. An opposite, recency kernel, where early evidence is under weighed, was observed in humans [12, 13]. Recently, in a large scale study of over 100 humans, we found that different people use different kernels with examples among the population of flat, primacy and recency effects. Intriguingly, however, the most popular kernel in our experiment was a ‘bump’ shaped kernel in which evidence in the middle of the stimulus is weighed more than either the beginning or the end [14].

In this work we show how dynamic divisive normalization [8] can act as a model for evidence accumulation in perceptual decision making. We provide theoretical results for how the model integrates evidence over time and show how dynamic divisive normalization can generate all of the four integration kernel shapes: primacy, recency, flat, and (most importantly) the bump kernel which is the main behavioral phenotype in our task [14]. In addition, we provide experimental evidence that divisive normalization can quantitatively account for human behavior in an auditory perceptual decision making task. Finally, with formal model comparison, we show that divisive normalization fits the data quantitatively as well as the state-of-the-art Drift Diffusion Model (DDM), the predominant model for perceptual evidence accumulation, with the same number of parameters.

## 2 Results

### 2.1 A divisive model of evidence accumulation

Our model architecture was inspired by the dynamic version of divisive normalization developed by Louie and colleagues to model neural activity during value based decision making [8]. We assume that the decision is made by comparing the activity in two pools of excitatory units, *R*_left_ and *R*_right_ (Figure 1a). These pools receive time varying input *C*_left_ and *C*_right_. In the Clicks Task (below), these inputs correspond to the left and right clicks, more generally they reflect the momentary evidence in favor of one choice over the other. An inhibitory gain control unit *G*, which is driven by the total activity in the excitatory network, divisively inhibits the *R* unit activity. The time varying dynamics of the model can be described by the following system of differential equations:

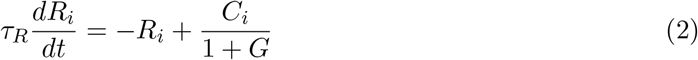

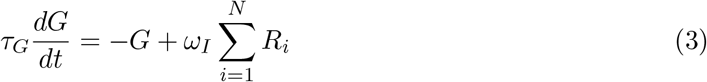

A decision is formed by comparing the difference in activity *δ* between the two *R* units

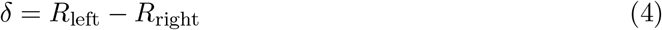

**Figure 1:**
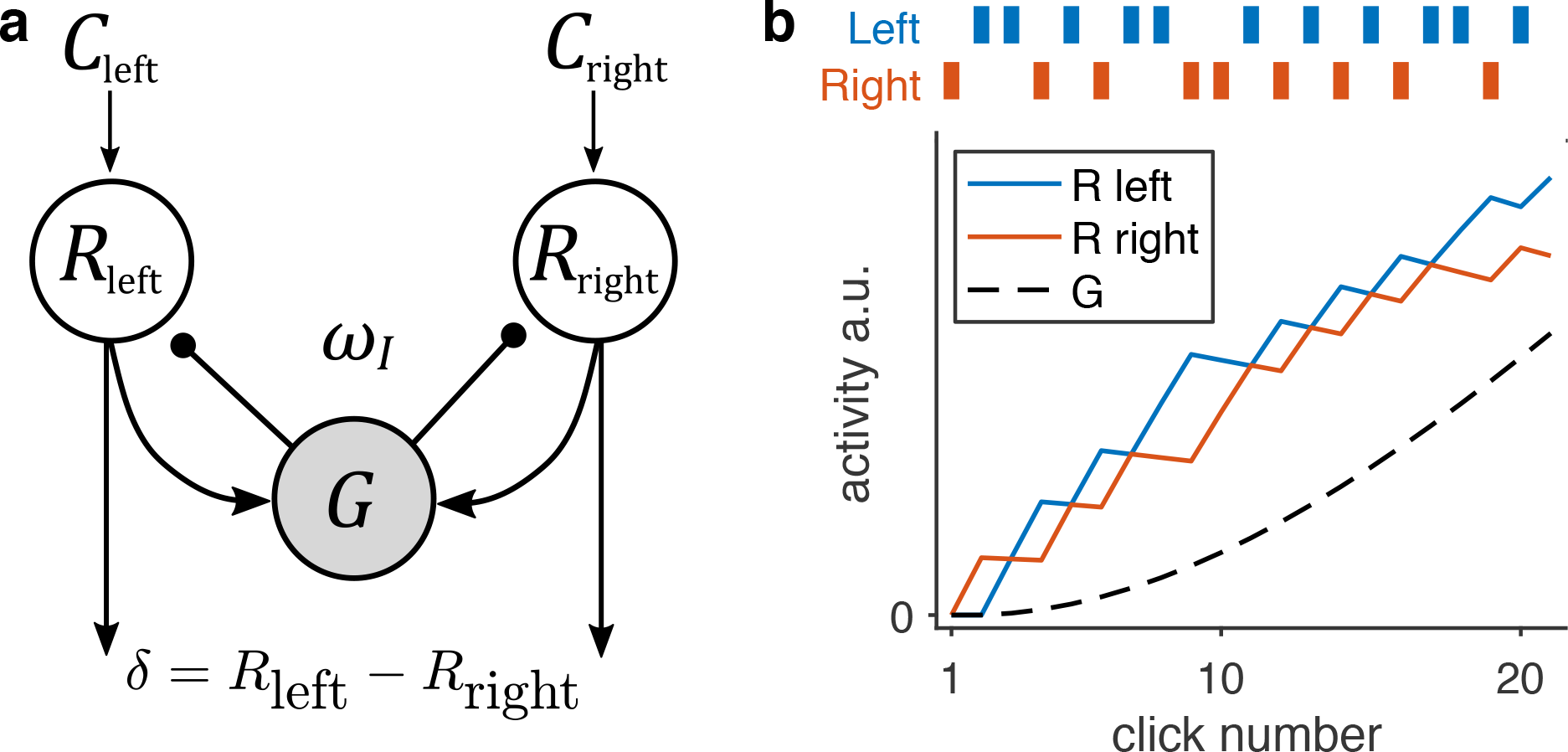
Dynamic divisive normalization schematic and simulated model dynamics. (a) Schematic of dynamic divisive normalization model. The two excitatory *R* units integrate punctate inputs *C* respective to left and right. The inhibitory *G* unit receives the sum of the two *R* unit activity weighted by *ω_I_*, and in turn divisively normalizes the input to *R*. (b) Results of the model activity simulated with *τ_R_* = 2.27, *τ_G_* = 11.10, and *ω_I_* = 36.20

Example simulated dynamics of the *R* and *G* units for punctate inputs (of the form used in the Clicks Task) are shown in Figure 1b. The model has three free parameters: *τ_R_*, *τ_G_*, and *ω_I_*. As is clear from this plot, the *R* unit activity integrates the input, *C*, over time, with each input increasing the corresponding *R* unit activity. In addition, closer inspection of Figure 1b reveals that the inputs have different effects on *R* over time — for example, compare the effect of the first input on the right, which increases *R*_right_ considerably, to that of the last input on the right, which increases *R*_right_ much less. This suggests that the model with these parameter settings integrates evidence over time, but with an uneven weighting for each input.

### 2.2 Dynamic divisive normalization generates different integration kernel shapes

How can we quantify the integration kernel — how much each piece of evidence weighs — given by a circuit that generates divisively normalized coding? We integrate the set of differential equations to provide an explicit expression for the integration kernel. We first consider the evolution of the difference in activity, *δ*, over time In particular, from equation (2) and (4), we can write

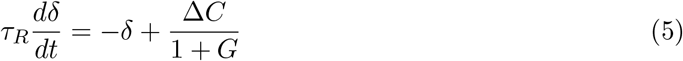

where ∆*C* is the difference in input,

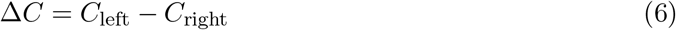

We can then integrate equation 5 using the ansatz

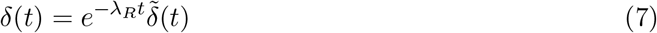

to compute the the following formal solution for *δ* as a function of time (for details of derivation see Methods Section 4.5):

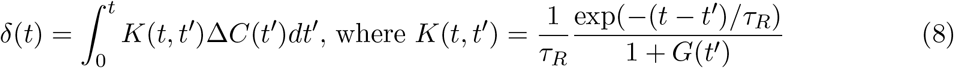

This expression shows explicitly that the activity of the network acts to integrate the inputs ∆*C* over time, weighing each input by the integration kernel function *K*(*t, t^′^*). Importantly, *K*(*t, t^′^*) represents the degree to which evidence ∆*C* at time *t^I^* contributes to the decision.

While clearly not a closed form expression for the integration kernel (notably *K*(*t, t^′^*) still depends on *G*(*t*)), equation (8) gives some intuition in how evidence is accumulated over time in this model. In particular, the kernel can be written as a product of two factors: an exponential function (Figure 2a left panel) and the inverse of the *G* activity (Figure 2a middle panel). The exponential function is increasing over time, and since *G* is increasing with time (Figure 1b)), the inverse of *G* is decreasing over time. Under the right conditions, the product of these increasing and decreasing functions can produce a bump shaped kernel, Figure 2a right panel.

**Figure 2:**
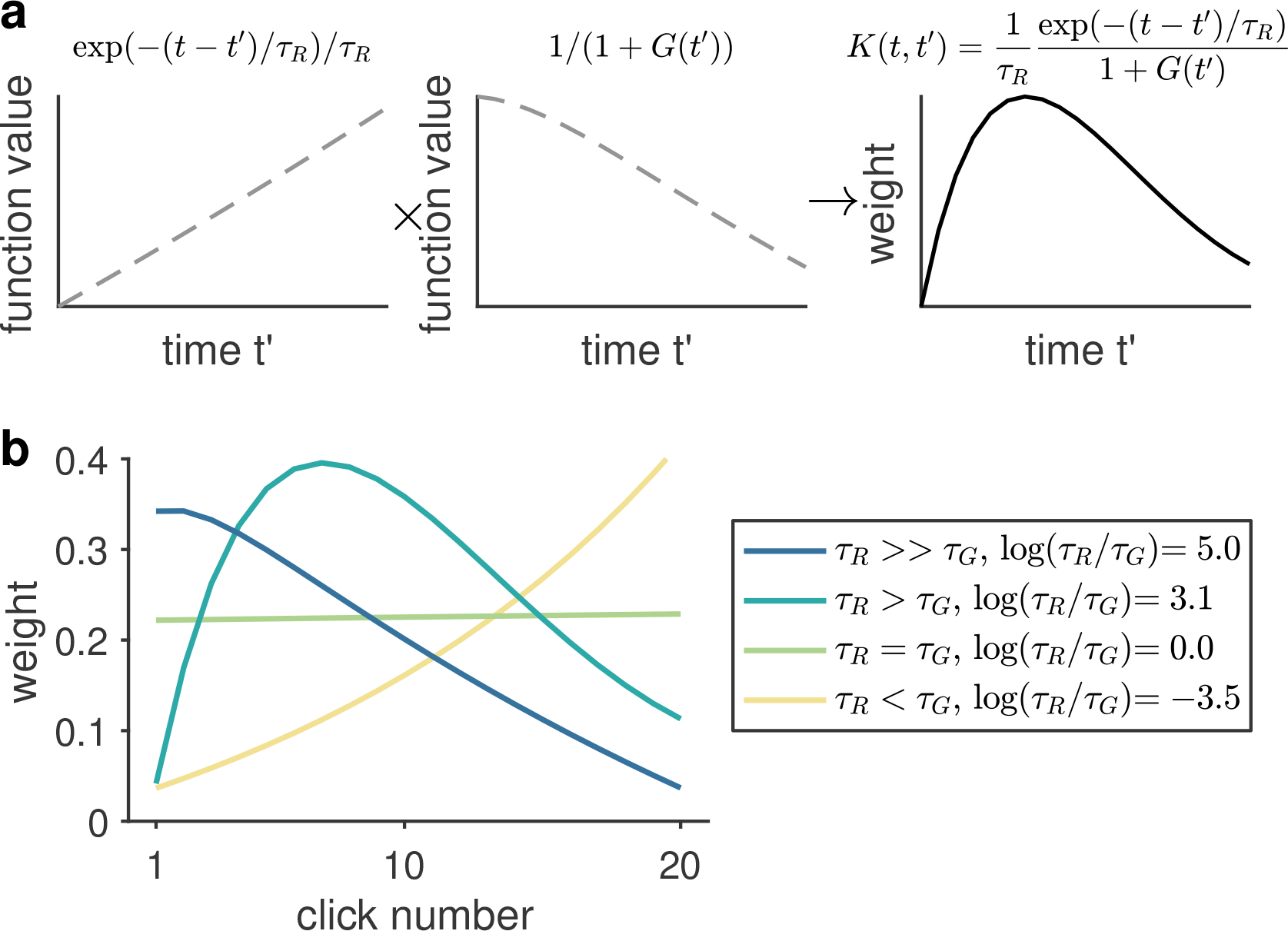
How divisive normalization generates different integration kernel shapes. (a) Example simulation demonstrates how the two components in the integration kernel *K* (equation (8) combines to generate a bump shaped kernel. *K* (right panel) is a product of an increasing exponential function (left panel) and the inverse of 1 + *G* (middle panel) which is decreasing over time. (b) Simulations of primacy, bump, flat, and recency integration kernels using decreasing log ratios of *τ_R_* and *τ_G_* to demonstrate that the shape of the integration kernel is determined by a balance between the rate of the leaky integration in *R* and the rate of the *G* inhibition.

More intuitively, we can consider integration kernel as being affected by two processes: the leaky integration in *R* and the increasing inhibition by *G*. If we consider the start of the train of clicks when *G* is small, the model acts as a leaky integrator (equation (2)), which creates a recency bias since earlier evidence is ‘forgotten’ through the leak. Over time, as *G* unit activity increases, *G* exerts an increasing inhibition on *R*, and when inhibition overcomes the leaky integration, later evidence was weighed less than the preceding evidence.

These intuitions suggest that the shape of the integration kernel is determined by a balance between how fast the leaky integration in *R* happens (the rate of *R*) and how fast the inhibitory *G* activity grows (the rate of *G*). These two rates are determined by the the inverse of the time constants *τ_R_* and *τ_G_* respectively — i.e. when *τ* is large, the rate is slow. The balance between the rate of *R* and the rate of *G* can then be described as the ratio *τ_R_/τ_G_* — i.e. when *τ_R_* is larger than *τ_G_*, *R* activity is slower than *G* activity, and similarly; when *τ_R_* is smaller than *τ_G_*, *R* activity is faster than *G* activity.

To investigate how integration kernels can change depending on a ratio between the rate of *R* and the rate of *G*, we simulated the integration kernel using different *τ_R_/τ_G_* ratios, and show that integration kernel shape changes from primacy, to bump, to flat, and then to recency as *τ_R_/τ_G_* decreases (Figure 2b). When *τ_R_/τ_G_* is much larger than 1, rate of integration is much slower than rate of inhibition by *G*. This inhibition suppresses input from later evidence, thus producing a primacy kernel. As *τ_R_/τ_G_* decreases towards one — *τ_R_* decreases and *τ_G_* increases, inhibition slows down and allows for leaky integration to happen, thus producing a bump kernel. When *τ_R_/τ_G_* reaches one, i.e. the two rates balances out, a flat kernel is generated. Finally, when *τ_R_/τ_G_* decreases to below one, leaky integration overcomes inhibition, generating a recency kernel.

### 2.3 Humans exhibit uneven integration kernels in a perceptual decision making task

To examine the model in the context of behaviour, we looked at behavioural data from 133 human participants. Most of this data (108 subjects) was previously published [14]. We observed that a large cohort of human participants weighed evidence unevenly when performing an auditory decision making task adapted from Poisson Clicks Task [10]. In this task, on every trial participants listened to a train of twenty clicks over one second at 20 Hz (Figure 3a). Each click was on either the left or the right side. At the end of the train of clicks participants decided which side had more clicks. Participants performed between 666 and 938 trials (mean 750.8) over the course of approximately one hour. Basic behaviour in this task was comparable to that in similar perceptual decision making tasks in previous studies [10, 11]. Choice exhibited a characteristic sigmoidal dependence on net difference in clicks between left and right (Figure 3b).

**Figure 3:**
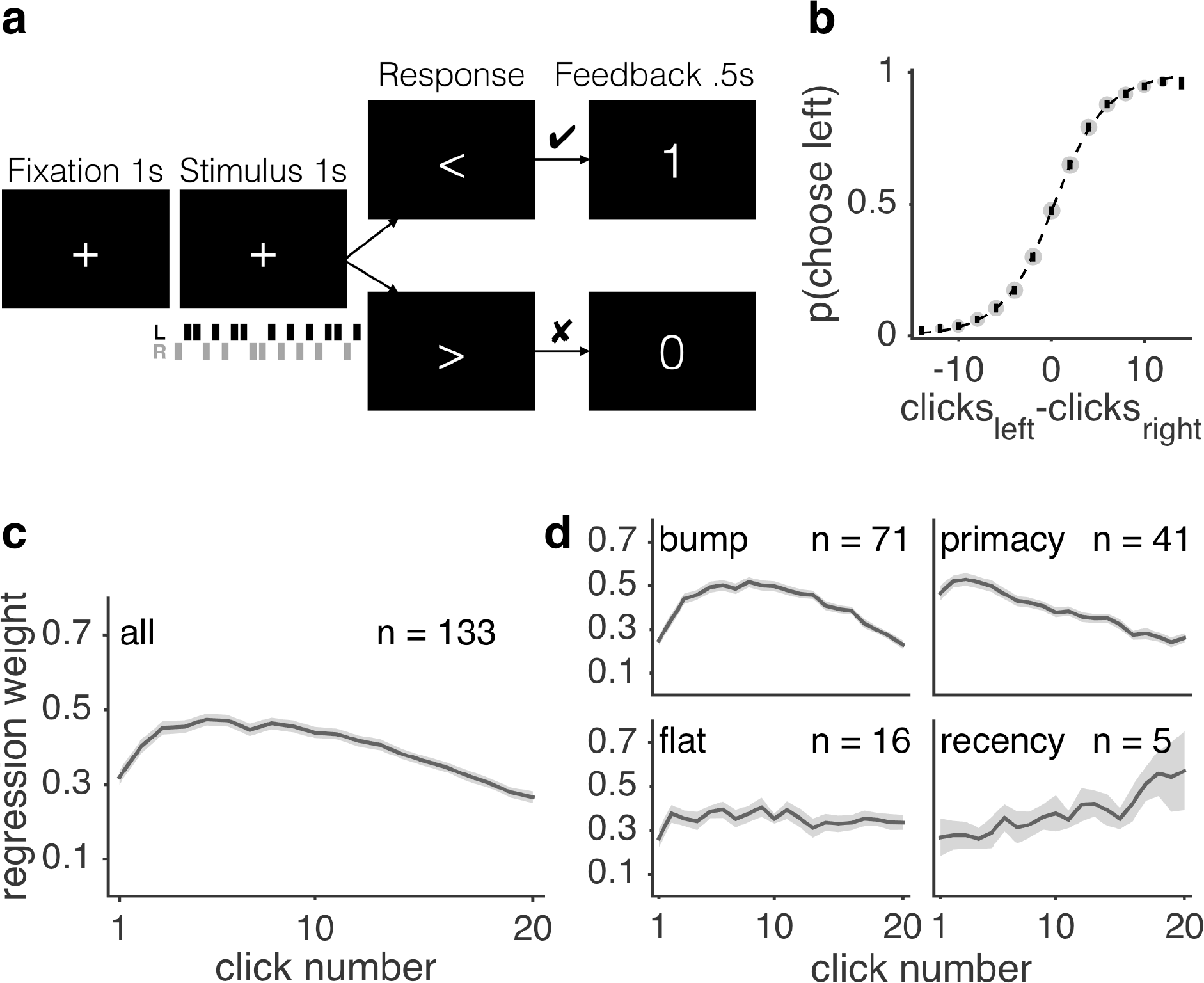
Humans exhibit uneven integration kernels in a perceptual decision making task. Task design, psychometric function, and different integration kernel shapes in human participants. Participants listened to a train of twenty clicks coming in either the left (L, black bars) or right (R, grey bars) ear for one second, and decided which side had more clicks. (b) Psychometric curve — choice probability (probability of choosing left) — showed sigmoidal relationship with difficulty (the difference in number of clicks between left and right). Error bars indicate s.e.m. across participants. Size of grey dots is proportional to number of trials. Dotted line indicates sigmoidal function fit. Shaded area indicate s.e.m. across participants. (c) Integration kernel, as 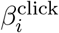, estimated from logistic regression (equation (9), averaged across all participants. (d) Plots of participants’ integration kernels grouped into four groups of different integration kernel shapes. All shaded areas indicate s.e.m. across participants.

We quantified the integration kernel, i.e. the impact of every click on choice, with logistic regression in which the probability of choosing left on trial *t* was given by

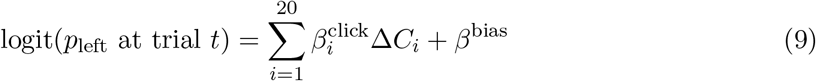

where ∆*C*_*i*_ was difference between left and right for the the *i*th click (i.e. ∆*C*_*i*_ = ∆*C*_*left,i*_ − ∆*C*_right*,i*_, therefore, ∆*C*_*i*_ was +1 for a left click and −1 for right). The integration kernel was quantified by the regression weights 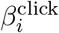, and *β*^bias^ characterized the overall bias.

We found that participants weighed the clicks unevenly over time (repeated measures ANOVA on 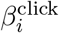: *F* (19, 2508) = 34.47, *p <* 0.00001). Importantly, post-hoc Tukey’s test showed that the middle of the kernel was significantly higher than either the beginning or the end of the click (3rd–9th clicks were higher than the 1st click, and 10th–12th clicks were higher than 16th–20th clicks, *p <* 0.00001), which indicated that on average participants tended to weigh the middle of the click train more than the beginning or the end, forming a ‘bump’ shaped kernel (Figure 3c). This uneven kernel shape contributed as a source of approximately 27% of the total errors in participants’ choices (see Supplementary Materials S1 and Figure S1).

To explore individual differences in integration kernels, we furthered quantified the shape of the integration kernel for each participant (for detailed description of categorization of integration kernels into shapes, see Supplementary Materials S2 and Figure S2). Specifically, we found that participants exhibited one of four distinct kernels: bump (n = 71, 53%), primacy (n = 41, 31%), flat (n = 16, 12%), and recency (n = 5, 4%) (Figure 3d).

### 2.4 Dynamic divisive normalization accounts for different integration kernels in human behavioural data

To investigate whether our divisive model could account for the range of integration kernels observed in human behavior, we fit the model to participants’ choices using a maximum likelihood approach. To fit the model to human behavior we assumed that a choice is made by comparing the activity in the two *R* units (i.e., *δ* = *R*_left_ *− R*_right_) with some noise, parameterized by *σ*, and an overall side bias (i.e. overall bias to either left or right). We also added an additional offset parameter *µ* to the kernel. With equation (8), the probability of choosing left is given by

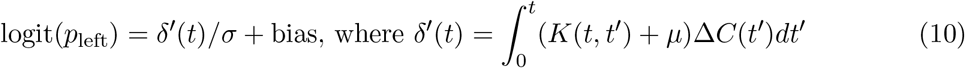

We computed the probability of a choice on a given trial at *t* = *T*, where *T* is the time at the end of the stimulus. The model has a total of six free parameters (*τ_R_*, *τ_G_*, inhibition weight *ω_I_*, noise *σ*, offset *µ*, and an overall bias). Using parameters that best fitted to each participant’s choice, we first reconstructed integration kernel from divisive normalization for each participant from the kernel function (equations (8) and (10)). Divisive normalization can account for all four types of integration kernel in human participants (Figure 4a and Supplementary Figure S3). We also used divisive normalization to generate simulated choices for each participant for each trial using the best fitting parameters, and showed that the resulting psychometric curve also matched well to that of human participants (Figure 4b and Supplementary Figure S4). The distribution of best fitted parameters is plotted in Supplementary Figure S5.

**Figure 4:**
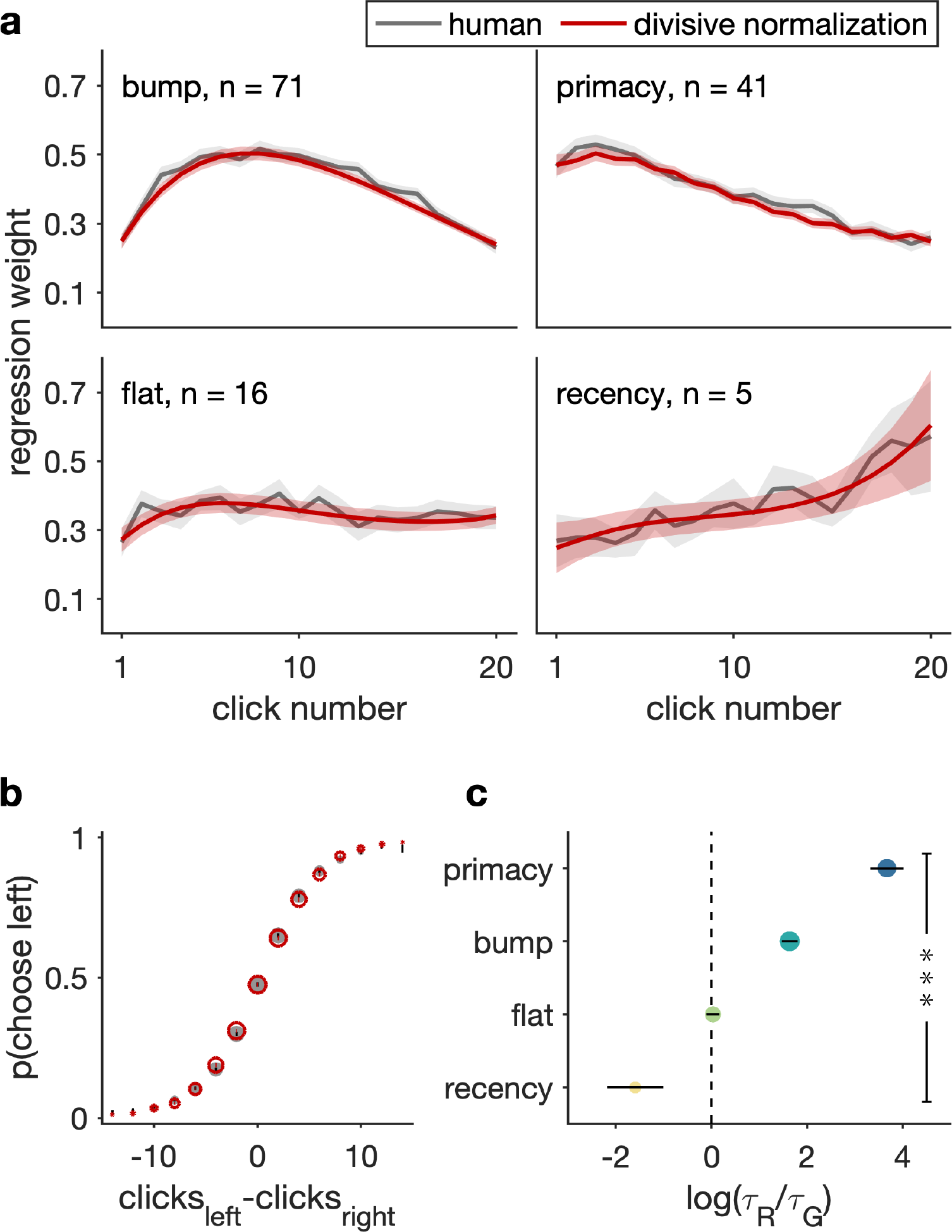
Dynamic divisive normalization accounts for integration kernels and psychometric curves in human participants. (a) Integration kernels generated by divisive normalization model fitted to human participants’ choices, compared to human integration kernels. Plots are grouped into groups of four different integration kernel shapes. Grey line indicates human integration kernels. Red line indicates model generated kernels. All shaded areas indicate s.e.m. across participants. (b) Psychometric curves generated by divisive normalization model compared to human psychometric curve. Grey circle indicates human psychometric curve. Red circle indicates model generated psychometric curve. Error bars indicate s.e.m. across participants. (c) Log ratios of fitted *τ_R_* and *τ_G_* values averaged within each integration kernel shape group. Log ratios of fitted *τ_R_* and *τ_G_* change significantly across kernel shape groups (***: Aone-way ANOVA *F* (3, 129) = 25.06, *p <* 0.001). Error bars indicate s.e.m. across participants within each group.

Our simulations in the previous section suggested that by shifting the balance between the integration and inhibition time constants (the ratio *τ_R_/τ_G_*), divisive normalization can generate the four types of kernel. We therefore examined the fitted parameter values in terms of *τ_R_/τ_G_*. As shown in Figure 4c log(*τ_R_/τ_G_*) is significantly different across kernel shapes (one-way ANOVA *F* (3, 129) = 25.06, *p* = 8 *×* 10^*−*13^). In particular, post-hoc Tukey test showed that log(*τ_R_/τ_G_*) in participants with bump, primacy, and flat kernels are significantly different from each other. Participants with bump or primacy kernels also have a significantly different log(*τ_R_/τ_G_*) from participants with recency kernel.

### 2.5 Dynamic divisive normalization performs as well as Drift Diffusion Model does in formal model comparison

Finally, to demonstrate that divisive normalization is comparable to an established model for such evidence accumulation tasks, we compared our model quantitatively with state-of-the-art Drift Diffusion Model (DDM) developed by Brunton, et al[10].

In its simplest form, the DDM assumes that an accumulator integrates incoming evidence over time (for example in our task the evidence +1 for a left click and −1 for a right click), with some amount of noise *σ_a_* added at every time step. In addition, a bias term is added to describe an overall bias to choosing either left or right. In an interrogation paradigm such as ours, a decision is made by comparing the accumulator activity with the bias when the stimulus ends, e.g. in our task, if the accumulator activity is larger than the bias, the model chooses left.

Later work added a ‘memory parameter’ *λ* to describe the extent to which the model is ‘forgetful,’ or ‘impulsive’ [15]. In particular a leaky accumulator (*λ <* 0) ‘forgets’ previous evidence and exhibits recency effect, while an impulsive accumulator (*λ >* 0) overweights early evidence and exhibits a primacy effect. When there is no memory noise (*λ* = 0), the integration kernel is flat.

Brunton and colleagues extended this standard DDM to include additional processes [10]: First, a bound, *B*, that describes the threshold of evidence at which the model makes a decision. In the context of an interrogation paradigm, evidence coming after the bound has been crossed is ignored. Second, a sensory adaptation process which controls the impact of successive clicks on the same size. This process is controlled by two adaptation parameters: 1) the direction of adaptation *φ*, which dictates whether the impact of a click on one side either increases (*φ >* 1) or decreases (*φ <* 1) with the number of clicks that were previously on the same side; and 2) a time constant *τ_φ_*, that determines how quickly the adapted impact recovers to 1.

Overall the Brunton model has six free parameters — neuronal noise, memory noise, bound, two parameters controlling sensory adaptation, and bias. We fit these parameters using the maximum likelihood procedure described in [10] and following code from [16]. We generated choices for each participant using the best fitting parameters, and computed an integration kernel for each participant using these model generated choices.

We found that divisive normalization can account for the behavioral data as well as the Brunton DDM can, both in formal model comparison using log likelihood, AIC, and BIC (Table 1), and in integration kernel and choice curve (Supplementary Materials S5 and Figure S7). Importantly, we show that the full Brunton DDM as reported in [10] with nine parameters accounts for the behavioral data equally well (Supplementary Materials S5 and Figure S8), suggesting that increasing the number of parameters did not improve model performance significantly. We also show that the standard form of DDM without bound or sensory adaptation does not account for participants’ choices as well as divisive normalization, even after accounting for the number of parameters with AIC and BIC (Supplementary Materials S5, Figure S6, and Table S1), suggesting that decreasing the number of parameters worsens the model performance. This result that divisive normalization can account for behavior as well as DDM can further supports divisive normalization as a model for evidence accumulation.

**Table 1:** Dynamic divisive normalization performs as well as Drift Diffusion Model does in formal model comparison. Comparison of log likelihood, AIC, and BIC scores between divisive normalization and DDM.

| model | log likelihood | AIC | BIC |
| --- | --- | --- | --- |
| Divisive normalization (6 param) | -357.8 | 727.5 | 755.2 |
| Brunton et al. DDM (6 param) [10] | -358.5 | 729.5 | 756.8 |

## 3 Discussion

In this work we proposed dynamic divisive normalization as a model for perceptual evidence accumulation. Theoretically, we provided a formal expression for the integration kernel — how this model weighs information over time — and how the shape of integration kernel is determined by the ratio of time constants within the model. Experimentally, we showed how dynamic divisive normalization can account for the integration kernels of human participants in an auditory perceptual decision making task. In addition, with quantitative model comparison, we show that dynamic divisive normalization explains participants’ choices as well as the state-of-the-art Drift Diffusion Model (DDM), the predominant model for such perceptual evidence accumulation tasks. Together, these results suggest that evidence accumulation can arise from a divisive normalization computation achieved through the interactions within a local circuit.

While our findings suggest that our model accounts well for human behavior in this one task, an obvious question is whether dynamic divisive normalization is at play in other types of evidence accumulation and in other decisions? For example, the drift diffusion model has been used to model evidence accumulation in a number of paradigms (from auditory clicks [10, 16, 17], to visual discrimination [18–20], to random dot motion [21–24]). Likewise the DDM can account for choice and reaction time data in quite different settings such as memory retrieval [25], cognitive control [26], and economic and value-based decision making [27–32]. Is divisive normalization also at play in these cases? If divisive normalization is a canonical neural computation, then the simple answer is ‘it must be,’ but whether its influence extends to behavior is largely unknown (although see the emerging literature on divisive normalization in economic and value-based decisions [6, 8, 33]).

If people are using divisive normalization in these decisions then what computational purpose does it serve? From a computational perspective, the DDM is grounded in the sequential probability sampling test (SPRT) which is the optimal solution to evidence accumulation problems for two-alternative decisions under certain assumptions [21, 34]. Is divisive normalization optimal under other decision making constraints? In this regard an intriguing finding by Tajima and colleagues suggest that divisive normalization may be almost optimal for multi-alternative decisions [35]. Other advantages of divisive normalization may be its ability to encode the state of the accumulator over a wide dynamic range of evidence [1, 36], or its relation to optimal Bayesian inference in some cases [37]. Of course an alternate account is that divisive normalization is necessary for other functions (e.g. balancing excitation and inhibition [1]) and the behavior we observe is simply the exhaust fumes of this function leaking out into behavior.

At the neural level, an obvious question is whether our neural model can explain neural data? In this regard it is notable that our model was adapted from Louie et al.’s model of lateral intraparietal (LIP) area neurons [8]. LIP has long been thought to contain a neural representation of the state of the accumulator [38–40] and it is likely that, just like Louie’s model accounts for the firing of LIP neurons in his task, our model may well be consistent with many of these past results. However, the accumulator account of LIP has recently been challenged [41–44] and other areas in prefrontal cortex [45–48] and striatum [49] have been implicated in evidence accumulation. Whether our divisive normalization explains neural firing in these areas is unknown.

Finally, we note that other neural network models of evidence integration have also been proposed, perhaps most importantly the model of Wang [50]. In its simplest form, the Wang model also considers two mutually inhibiting units that, superficially, look similar to the *R* units in Figure 1. However, the dynamics of the Wang model and the way it makes decisions are quite different. In particular, the mutual inhibition is calibrated in such a way that the Wang network has two stable attractor states corresponding to the outputs of the decision (e.g. left or right). The input, combined with the dynamics of the network, pushes the network into one of the two attractor states, which corresponds to the decision the network makes. Because the attraction of an attractor gets stronger the closer the network gets to it, the initial input to the model has a strong effect on the ultimate decision leading to a pronounced primacy effect in the Wang model. In contrast to Wang attractor model, our dynamic divisive normalization is essentially a line attractor network, with a single fixed point in *A*-*G* space which is stable for all values of *δ* (Supplementary Materials S6). This structure allows divisive normalization to exhibit a number of different integration kernels as shown in Figure 2 depending on the parameters.

In sum, dynamic divisive normalization can account for human behavior in an auditory perceptual decision making task, but much evidence remains to be accumulated before we can be sure that this model is correct!

## 4 Methods

### 4.1 Participants

188 healthy participants (University of Arizona) took part in the experiment. We analyzed the data from 133 participants (55 participants were excluded due to poor performance — accuracy lower than 60%). All human participants provided informed written consent prior to the experiment, and both experiments were approved by the local ethics committee.

### 4.2 Experimental procedures

Participants made a series of auditory perceptual decisions. On each trial they listened to a series of 20 auditory “clicks” presented over the course of 1 second. Clicks could be either ‘Left’ or ‘Right’ clicks, presented in the left or right ear. Participants decided which ear received the most clicks. In contrast to the Poisson Clicks Task [10], in which the click timing was random, clicks in our task were presented every 50 ms with a fixed probability (*p* = 0.55) of occurring in the ‘correct’ ear. The correct side was determined with a fixed 50% probability.

Participants performed the task on a desktop computer, while wearing headphones, and were positioned in chin rests to facilitate eye-tracking and pupillometry. They were instructed to fixate on a symbol displayed in the center of the screen, where response and outcome feedback was also displayed during trials, and made responses using a standard keyboard. Participants played until they made 500 correct responses or 50 minutes of total experiment time was reached.

### 4.3 Psychometric curve

Psychometric curves show the probability of the participant responding leftward as a function of the difference between the number of left clicks and the number of right clicks *C_left_ − C_right_*. The identical procedure was used to produce model-predicted curves, where the model-predicted probability of choice on each trial was used instead of the participants’ responses.

### 4.4 Integration kernel

To measure the contribution of each click to the participant’s choice on each trial (Figure 2A), we used logistic regression given by logit(*Y*) = *βX*, where *Y ∈ {*0, 1*}* is a vector of the choice on each trial and *X* is a matrix in which each row is the twenty clicks (∆*C* = *C*_left_ *− C*_right_) on that trial, coded as +1 for left and −1 for right. The identical procedure was used to produce model-predicted integration kernels, where the model-predicted choice on each trial was used instead of the participants’ responses.

### 4.5 Derivation of kernel function of divisive normalization

The model and the dynamical equations for *R* and *G* are described in the main text. These are reproduced here:

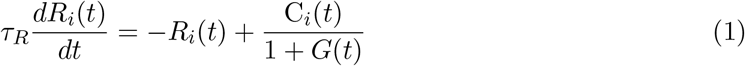

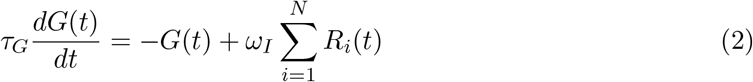

From equation (1) we can consider how the difference in activity *δ*(*t*) = *R*_left_(*t*) *− R*_right_(*t*) changes over time:

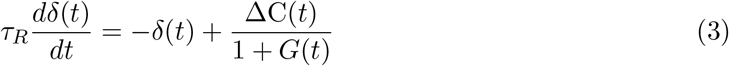

where ∆*C*(*t*) = *C*_left_(*t*) *− C*_right_(*t*) describes the difference in input over time.

To derive a formal expression for the kernel function, we integrate equation (3) using the ansatz:

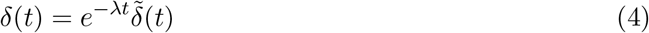

Taking the derivative of (4) and multiplying both sides with *τ_R_*, we get:

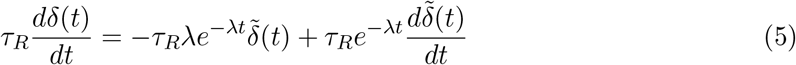

Combining equations (3), (4), and (5), we get:

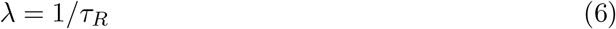

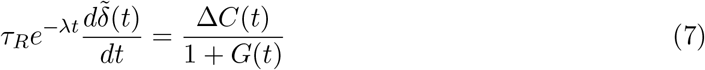

Integrating equation (7) we get:

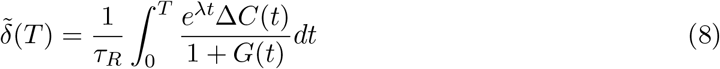

Substituting equation (8) back into equation (4), we get

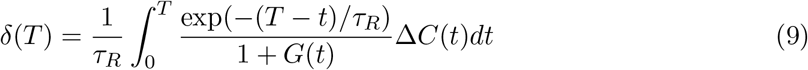

## Acknowledgments

We thank Samuel Feng for his advice and discussion. Maxwell Alberhasky, Chrysta Andrade, Daniel Carrera, Kathryn Chung, Michael de Leon, Zamigul Dzhalilova, Asha Esprit, Abigail Foley, Emily Giron, Brittney Gonzalez, Anthony Haddad, Leah Hall, Maura Higgs, Marcus Jacobs, Min-Hee Kang, Kathryn Kellohen, Neha Kwatra, Hannah Kyllo, Alex Lawwill, Stephanie Low, Colin Lynch, Alondra Ornelas, Genevieve Patterson, Filipa Santos, Shlishaa Savita, Catie Sikora, Vera Thornton, Guillermo Vargas, Christina West, and Callie Wong for help in running the experiments.

## Author contributions

W. K. analyzed the data. W. K. and R. C. W. did the modelling work. T. H. collected the data. T. H. and R. C. W. designed the experiment. W. K. and R. C. W. wrote the manuscript. All three authors contributed to interpretation of the results and critical discussion.

## Declaration of Interests

The authors declare no competing financial or non-financial interests as defined by Nature Research.

## Data availability

The data sets generated and analysed during the current study are available from the corresponding author upon reasonable request.

## Code availability

Experiment code was created with Psychtoolbox-3 and custom MATLAB code. All analyses were created with custom MATLAB and R code. All code will be uploaded to https://github.com/janekeung129/click-model

## Supplementary Materials

### S1 Errors contributed by uneven integration kernel

To understand how much error an uneven integration kernel introduces, we estimated error rates using simulations that keep the uneven integration kernel as the only source of error.

We first used logistic regression to estimate the regression weights of each click for each participant, as described in Main Text equation (9), with the equation replicated here:

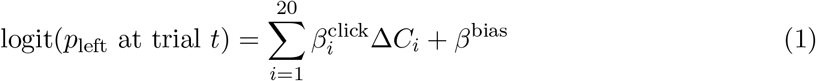

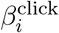 is the weight of the *i*th click on choice, and *β*^bias^ is the weight of an overall side bias (i.e. the weight of always choosing left).

We then simulated participants’ choices by reversing the logistic regression — for each participant, using the estimated betas, we computed *p*_left_, the probability of that participant choosing left on a given trial, using the same equation (equation (1)). We then reproduced the participant’s choices by randomly drawing from a binomial distribution with the computed *p*_left_ for each trial.

Then we compared the error rate of the simulated choices to participants’ actual choices. We showed in the rightmost two data points in Figure S1 that we reproduced the same total error rate in simulations using the original estimated regression weights (20.6%) as the total error rate in human participants’ data (20.2%).

We then removed the overall side bias by setting *β*^side^ to be zero. We also removed overall noise by making the choices deterministic. That is, instead of randomly drawing from a binomial distribution using *p*_left_, we asked whether logit(*p*_left_) at a given trial is larger or smaller than zero: if logit(*p*_left_) is larger than zero, choose left, and if logit(*p*_left_) is smaller than zero, choose right. If logit(*p*_left_) is equal to zero, then flip a coin with 50% probability. By removing these other sources of errors, we asked how much errors the uneven integration kernel shape contributes to. We showed that an uneven integration kernel shape contributed to 5.5% error rate (Figure S1), which accounted for 27.2% of the total error rate. We also showed that by removing the uneven integration kernel, simulations showed zero error rates.

**Figure S1:**
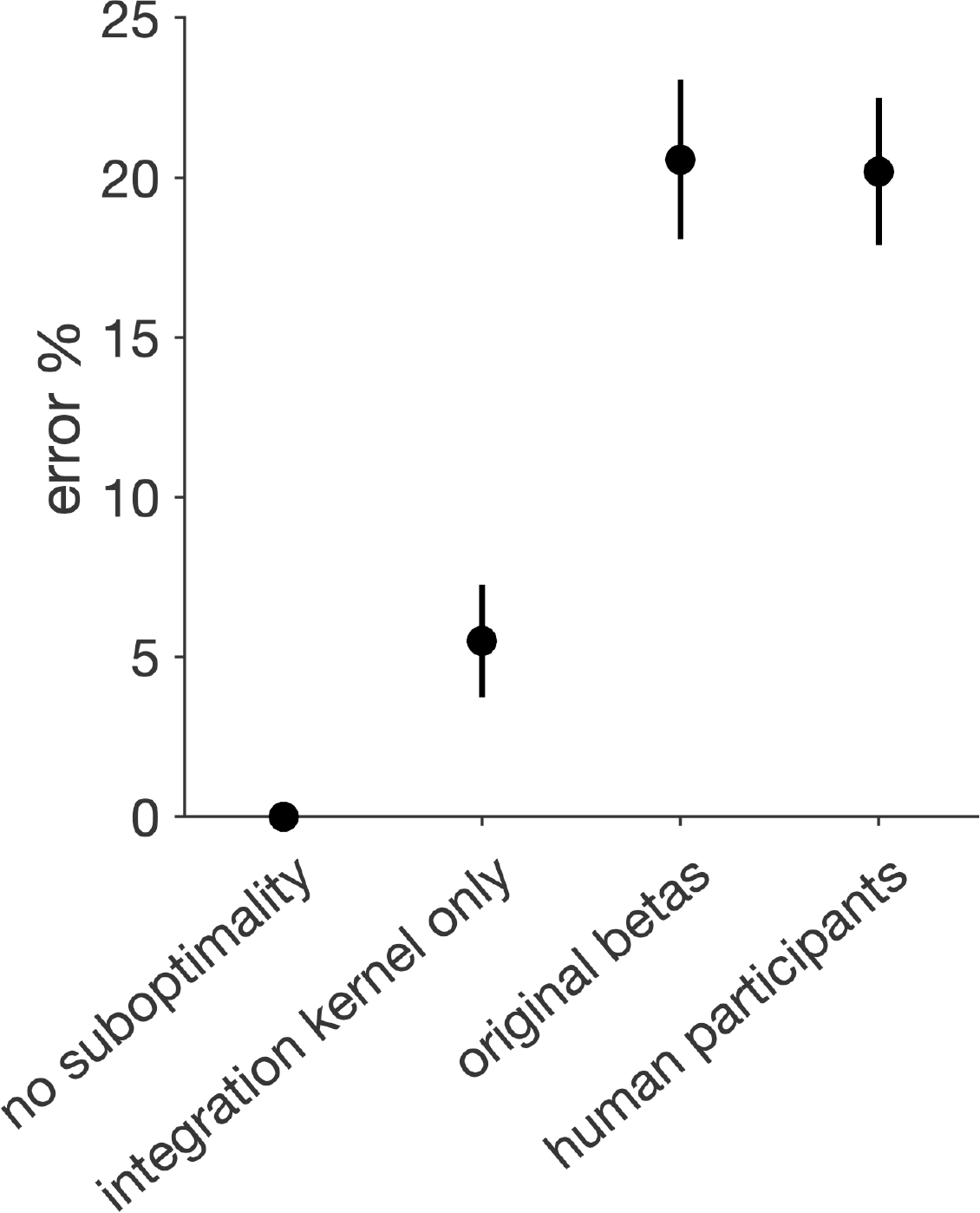
Error rates from simulated choices compared to human participants. From left to right: error rates of simulated choices with 1) no suboptimalities, 2) only uneven integration kernel, and 3) original betas, and error rate of human participants.

### S2 Categorizing integration kernels into shapes

To categorize the kernel for each participant into one of the four shapes, we fit polynomial functions with different degrees to participants’ choices, and selected the best fitting model with model comparison using the Akaike Information Criterion (AIC) to account for the different number of free parameters. In particular, we assume that the probability of choosing left at trial *t* is (the logit of) the weighted sum of clicks, where the weights are from a polynomial function, as shown in the following equation:

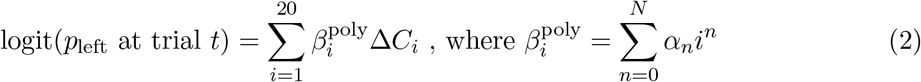

We fitted three different polynomial functions by changing N from 0 to 2: constant, linear, and quadratic. We then selected the best fitting function for each participant by comparing the fits from different polynomials with AIC. We categorized each participant’s integration kernel into one of the four shapes using the following criteria: (1) flat: kernel was best fit with the constant function; (2) primacy: kernel was best fit with linear function with a negative slope (*α*_1_), or with quadratic function with a minimum (*α*_2_ *>* 0) and the minimum is located later than the 10th click, or with quadratic function with a maximum (*α*_2_ *<* 0) and the maximum is located earlier than the 2nd click; (3) recency: kernel was best fit with linear function with a positive slope, or with quadratic function with a minimum (*α*_2_ *>* 0) and the minimum is located earlier than the 10th click, or with quadratic function with a maximum (*α*_2_ *<* 0) and the maximum is located later than the 18th click; (4) bump: kernel that did not meet the previous three criteria (i.e. kernel was best fit with quadratic function and was neither primacy nor recency). Individual integration kernels and their categorizations are plotted in Figure S2.

**Figure S2:**
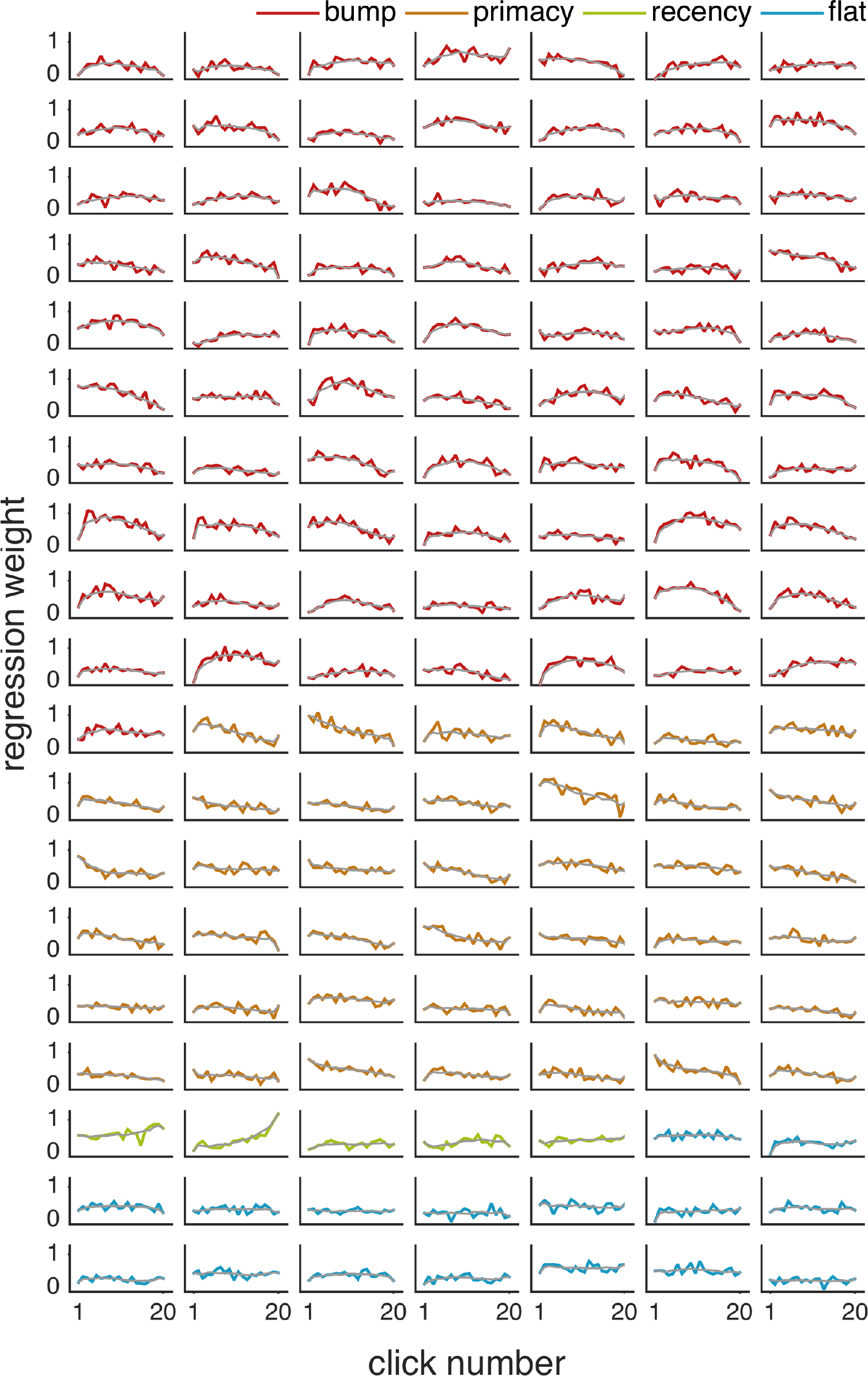
Individual integration kernel plots. Colored lines are regression weights of clicks from equation (9). Plots are sorted and color coded by kernel shape. Light grey line shows smoothed integration kernel.

### S3 Individual plots of integration kernels and psychometric curves generated by divisive normalization

**Figure S3:**
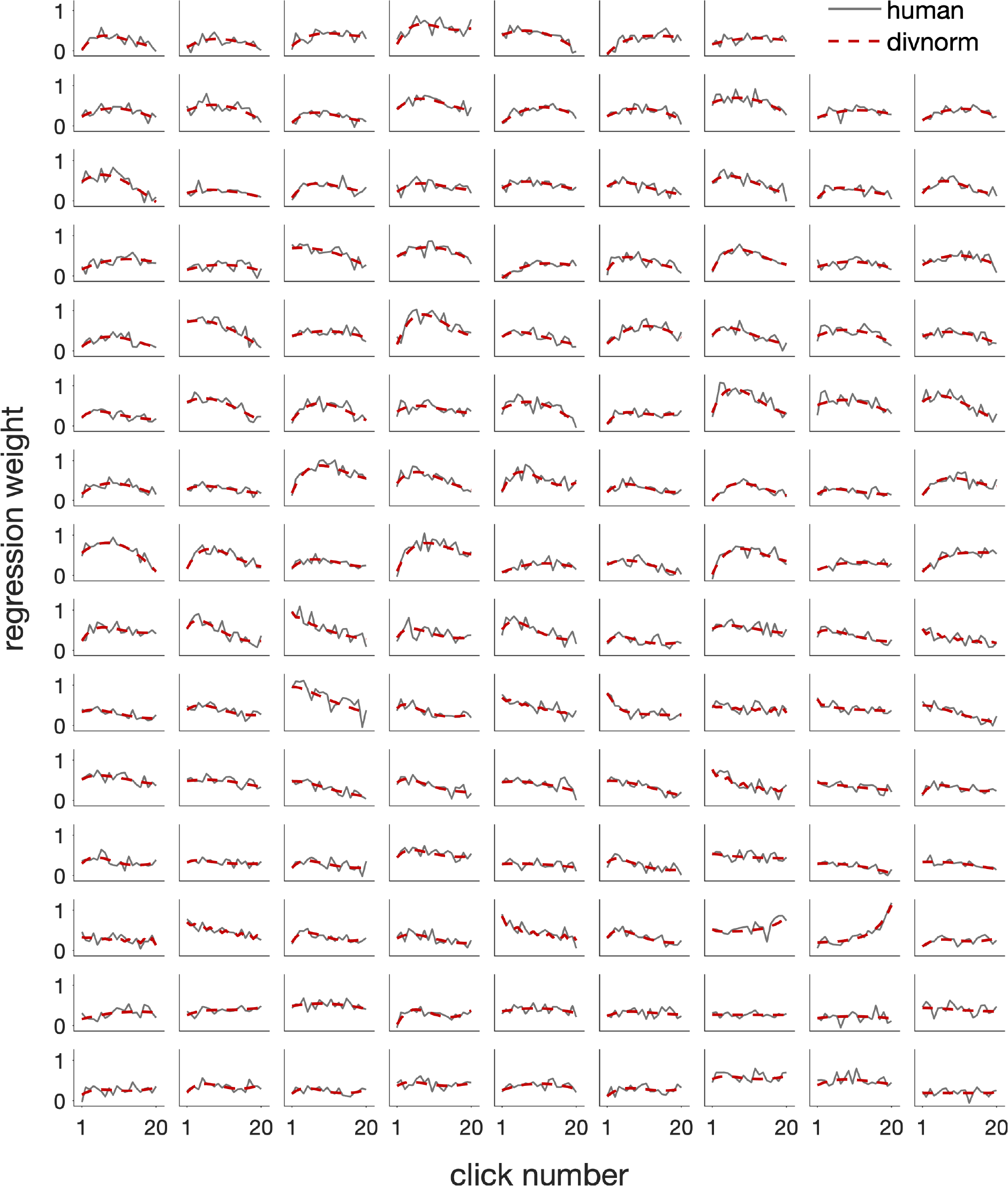
Individual integration kernels generated by divisive normalization compared to human integration kernels.

**Figure S4:**
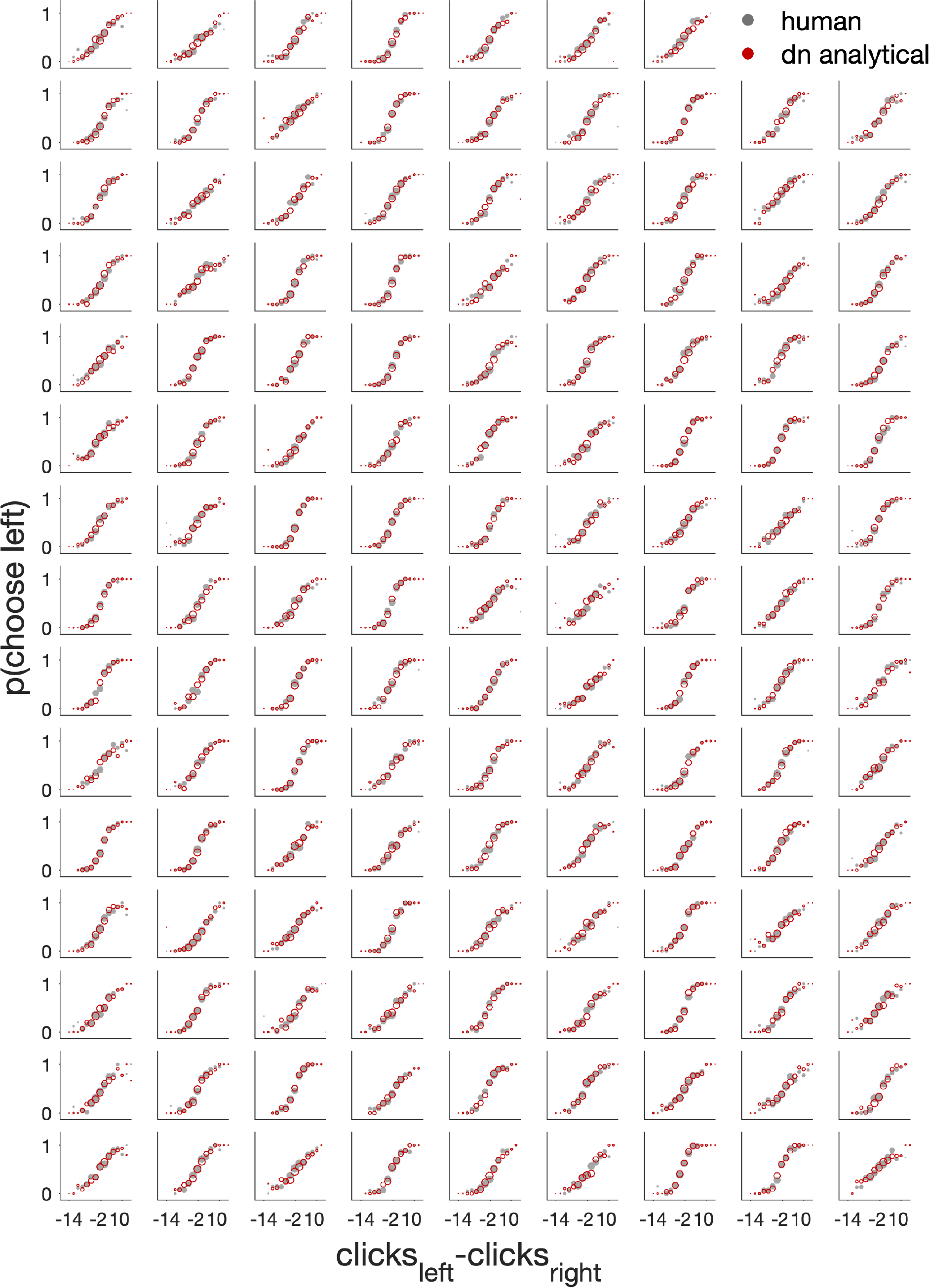
Individual psychometric curves generated by divisive normalization compared to human psychometric curves.

### S4 Histogram of fitted divisive normalization parameters

**Figure S5:**
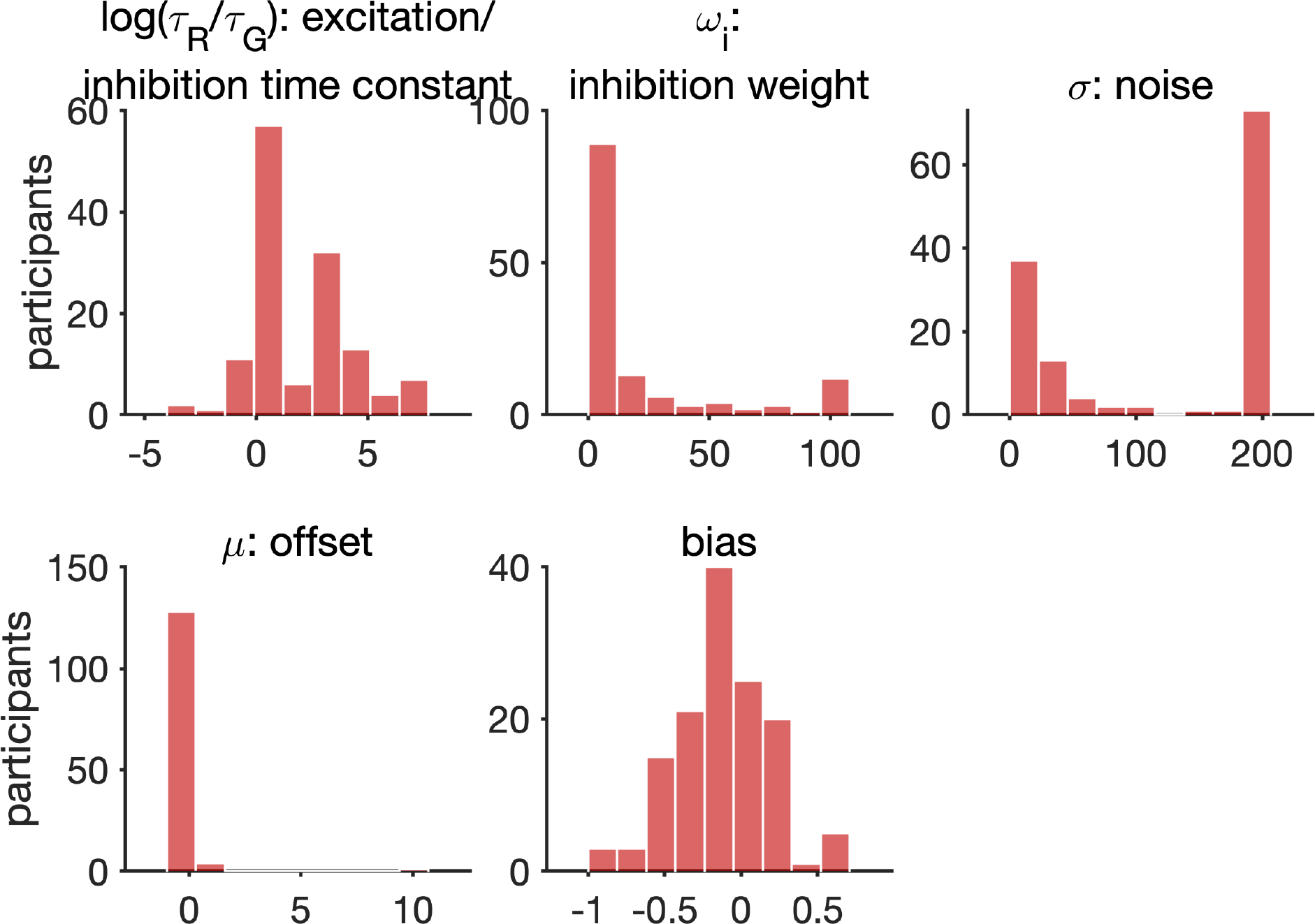
Histogram of fitted divisive normalization parameters.

### S5 DDM

#### S5.1 Standard DDM does not fit bump kernel

We first tested how standard DDM fits to the behavioural data. Intuitively, the standard DDM (i.e. without a bound) should not be able to generate a ‘bump’ shaped integration kernel. The most standard form of DDM:

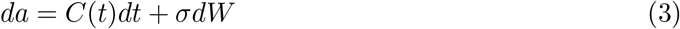

with only drift (input, i.e. clicks *C*) and diffusion (noise added by Wiener process *W*), and without any bound, would predict that every piece of evidence over time is integrated with equal weight — i.e. a flat integration kernel. An extension can be added to the standard DDM in the form of a ‘memory noise’ to account for primacy or recency integration kernels as well. This ‘memory’ parameter *λ* arises out of leaky competitive accumulators (LCA) model under certain constraints [15, 21]:

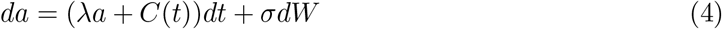

*λ* acts to maintain the memory of the evidence estimate. When memory is subtractive (*λ <* 0), DDM becomes leaky and earlier evidence is ‘forgotten’ and thus weighed less, creating a recency bias. When memory is additive (*λ >* 0), accumulator activity drifts exponentially over time, and the direction of the drift is determined by the initial stimulus, thus creating a primacy effect. When *λ* = 0, LCA (equation (4)) reduces to standard DDM (equation (3)).

We fitted the standard DDM without bound to participants’ choices using a maximum likelihood approach. An analytical solution of the probability of choosing a certain side exists for DDM without bound under fixed decision time protocol [22]. Specifically, assuming the probability distribution of the initial accumulator state is Gaussian (with initial mean *µ*_0_ and initial variance *v*_0_), the probability distribution of the accumulator state at the end of the stimulus train is also Gaussian, and the mean and variance can be computed analytically. Thus, the probability of choosing one side is the cumulative normal distribution.

We used maximum likelihood approach to fit these five parameters in LCA: initial mean *µ*_0_, initial variance *v*_0_, memory noise *λ*, noise *σ*, and an overall bias. We generated choices from the model using best fitting parameters. Integration kernels from DDM can be reconstructed by regressing the stimulus onto model generated choices using logistic regression (equation (9)).

Confirming our intuition, we show that the standard DDM can produce a primacy, recency, and flat effect, but cannot fit to the bump kernel (Figure **??**a).

**Figure S6:**
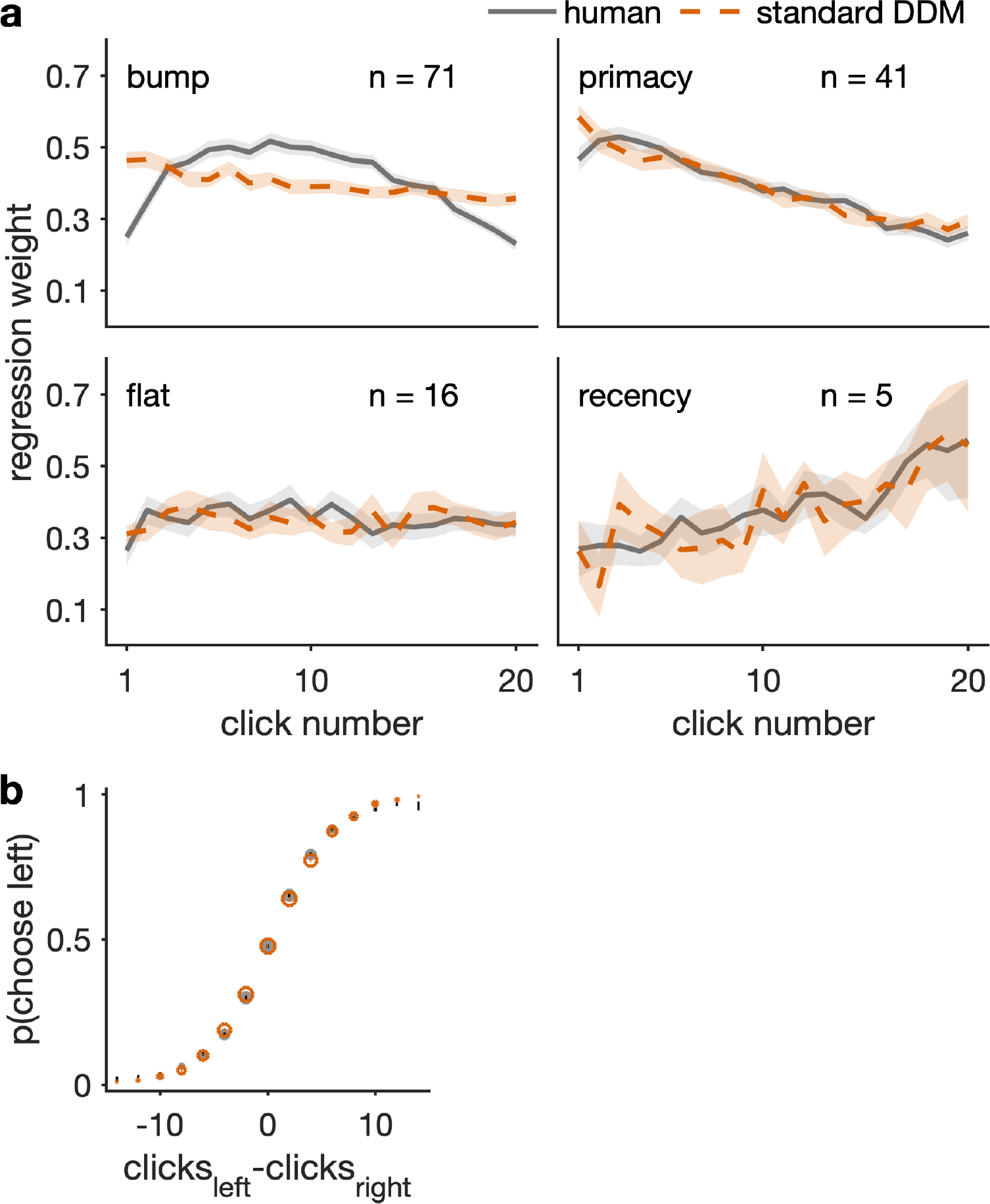
Standard DDM fits

#### S5.2 6 parameter Brunton et al. DDM fits participants’ behaviour

We then fitted the Brunton et al. DDM [10] to the behavioural data. The key differences between the Brunton et al. DDM and standard LCA model is the addition of two processes. The first is a ‘sticky’ bound — that is, a decision is made either at the end of the stimulus train, or at the time when the accumulator hits the bound, depending on which event happens earlier. The second is a sensory adaptation process which controls the actual impact of a click (without adaptation the impact of every click will always be 1). The process is controlled by two parameters: 1) the direction of adaptation — either no adaptation, depression (the impact of each click is smaller than 1, or facilitation (the impact is larger than 1), and 2) a time constant that determines how quickly the adapted impact recovers to 1.

**Figure S7:**
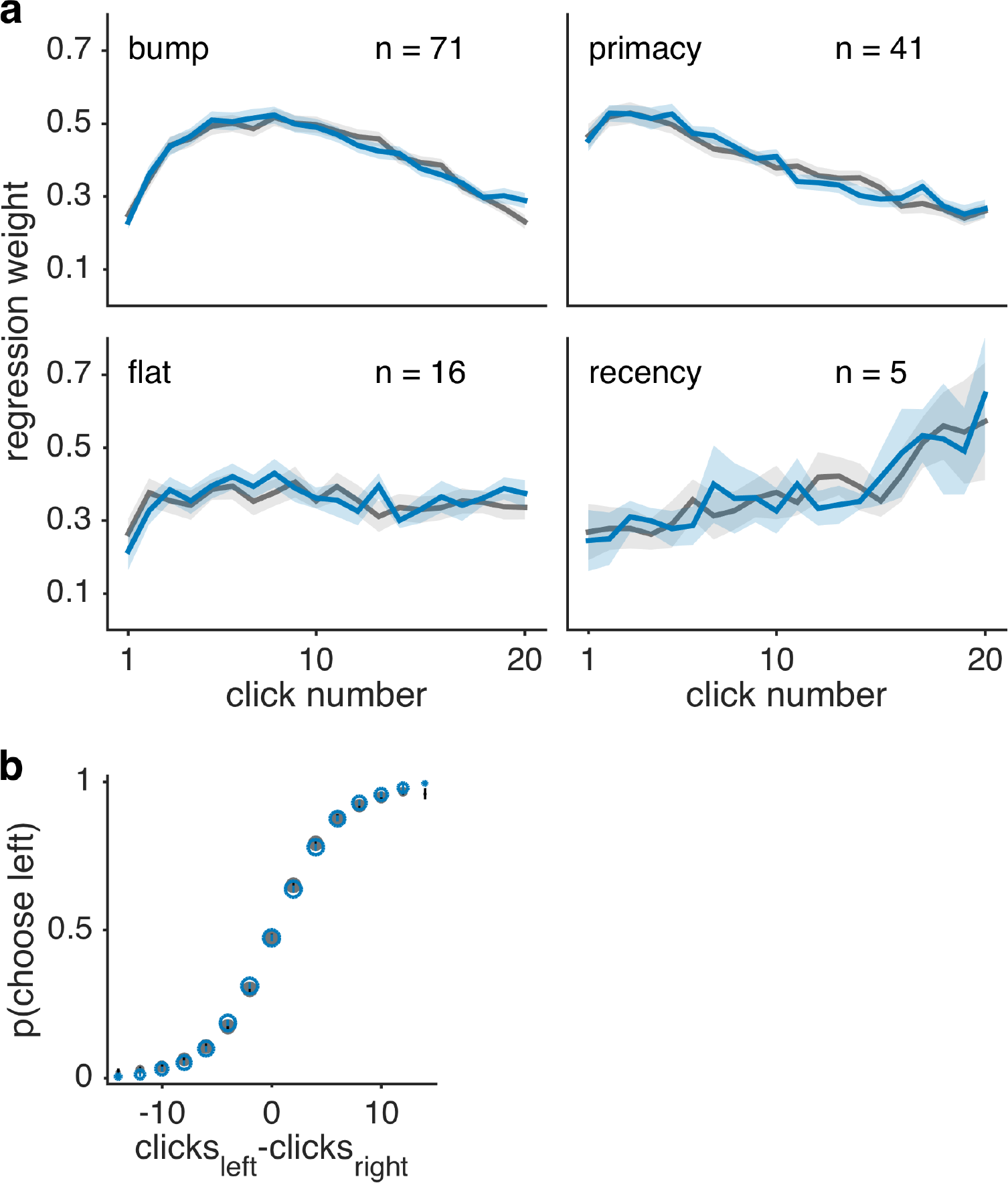
Brunton 6 param DDM fits.

To fit the Brunton el al. model, the probability distribution of accumulator state evolves over time following a similar logic to the standard DDM. To model the bound in the probability distribution space, a ‘sticky’ bound is added to DDM such that when the probability mass hits the bound, it sticks to the bound. The evolution of the probability distribution for each trial has to be computed numerically, until the end of the stimulus. Similar to the case of standard DDM, the probability of choosing one side is the cumulative distribution of the probability distribution at the end of the stimulus train. This cumulative distribution is also computed numerically. We fitted the model to participants’ choices using code provided in [16]. We show that Brunton DDM fits to all four integration kernel shapes.

#### S5.3 9 parameter Brunton et al. DDM fits behaviour as well as 6 param does

The original model reported in Brunton et al. [10] has three additional parameters: 1) an additional noise parameter *σ_s_* that characterizes the noise added at each incoming stimulus (i.e. click), 2) a noise parameter *σ_i_* that characterizes the amount of noise in the initial state of the accumulator, and 3) a lapse rate that characterizes the probability of a random response being made. We fitted the Brunton et al. DDM with the nine parameters to our behavioural data and show that the DDM fit to the data as well as the DDM with six parameters.

**Figure S8:**
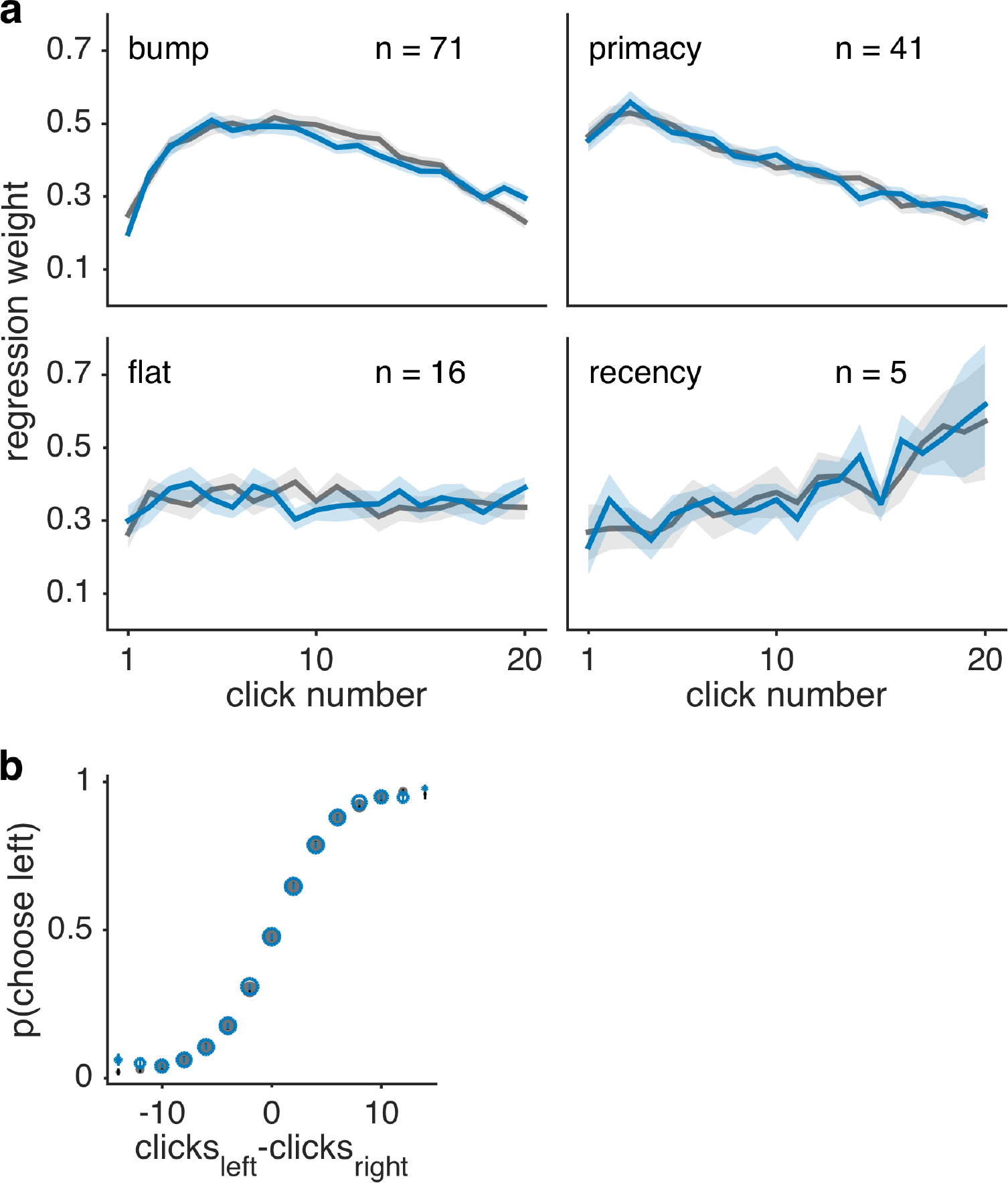
Brunton 9 param DDM fits.

#### S5.4 Formal model comparison

**Table S1:** scores

| model | log likelihood | AIC | BIC |
| --- | --- | --- | --- |
| Divisive normalization (6 param) | -357.8 | 727.5 | 755.2 |
| Brunton et al. DDM (6 param) [10] | -358.5 | 729.5 | 756.8 |
| Brunton et al. DDM (9 param) [10] | -357.7 | 733.1 | 774.7 |
| Standard DDM [22] | -364.2 | 738.5 | 761.6 |

#### S5.5 Histogram of fitted parameters of 6 param Brunton DDM

**Figure S9:**
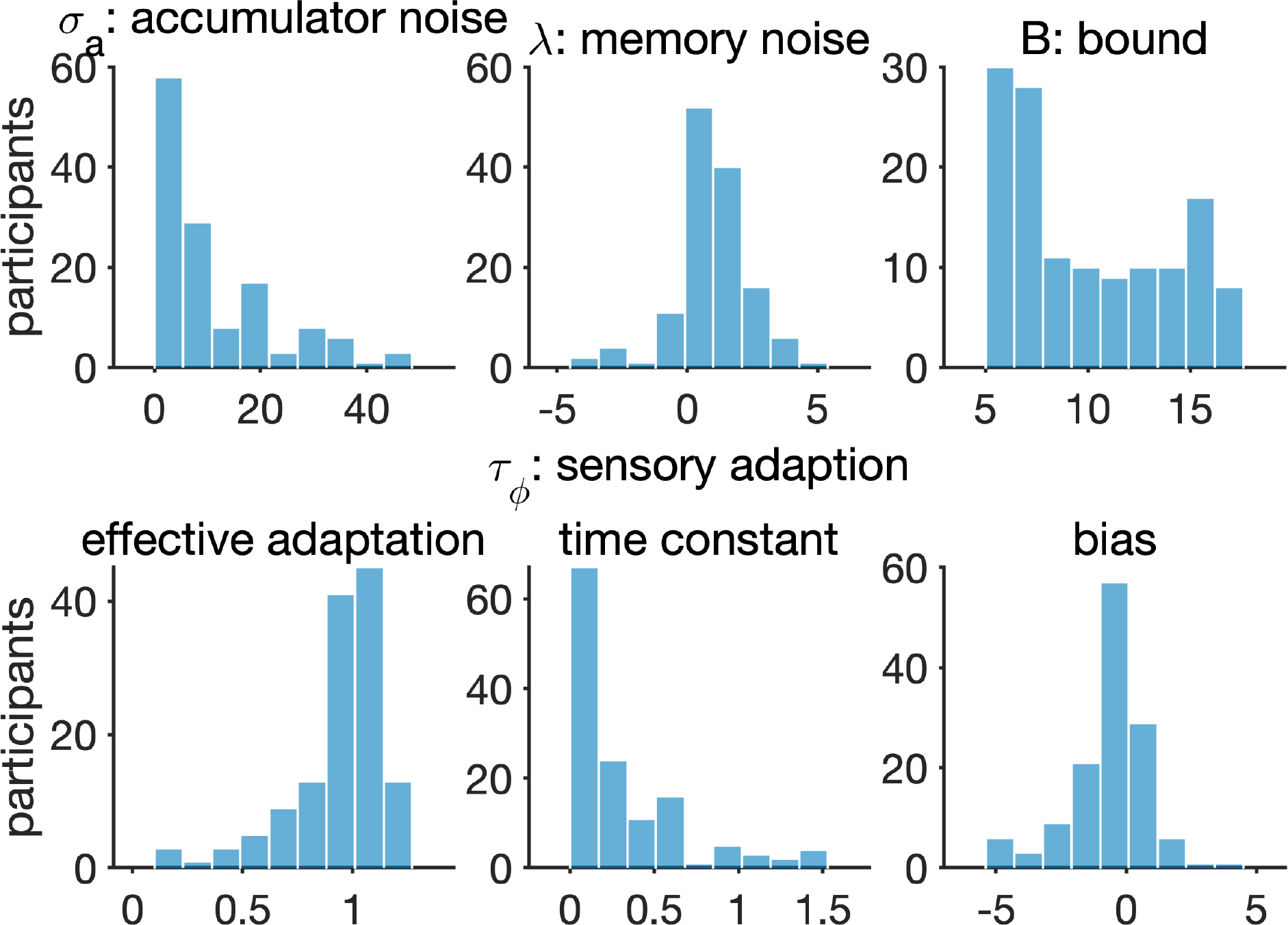
Histogram of fitted DDM parameters.

### S6 Attractor space of divisive normalization

We show that there is one attractor in the *A*-*G* space where *A* is the total network activity:

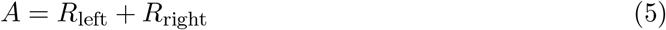

Since with our experimental design, there is always a click input at each time point (so either *C*_left_ = 1 or *C*_right_ = 1), the sum of inputs into the network is constant at 1 over time. Thus, the set of differential equations for the *A*-*G* space are:

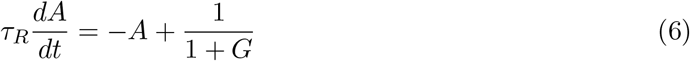

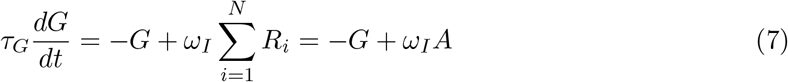

Analytically, a full proof for the existence of a unique equilibrium point that is asymptotically stable for general families of this set of differential equations was provided by Louie, et al. [8]. Numerically, we produce plots of vector field and model trajectory to demonstrate the stable equilibrium point for divisive normalization in the *A*-*G* space (Figure S10).

**Figure S10:**
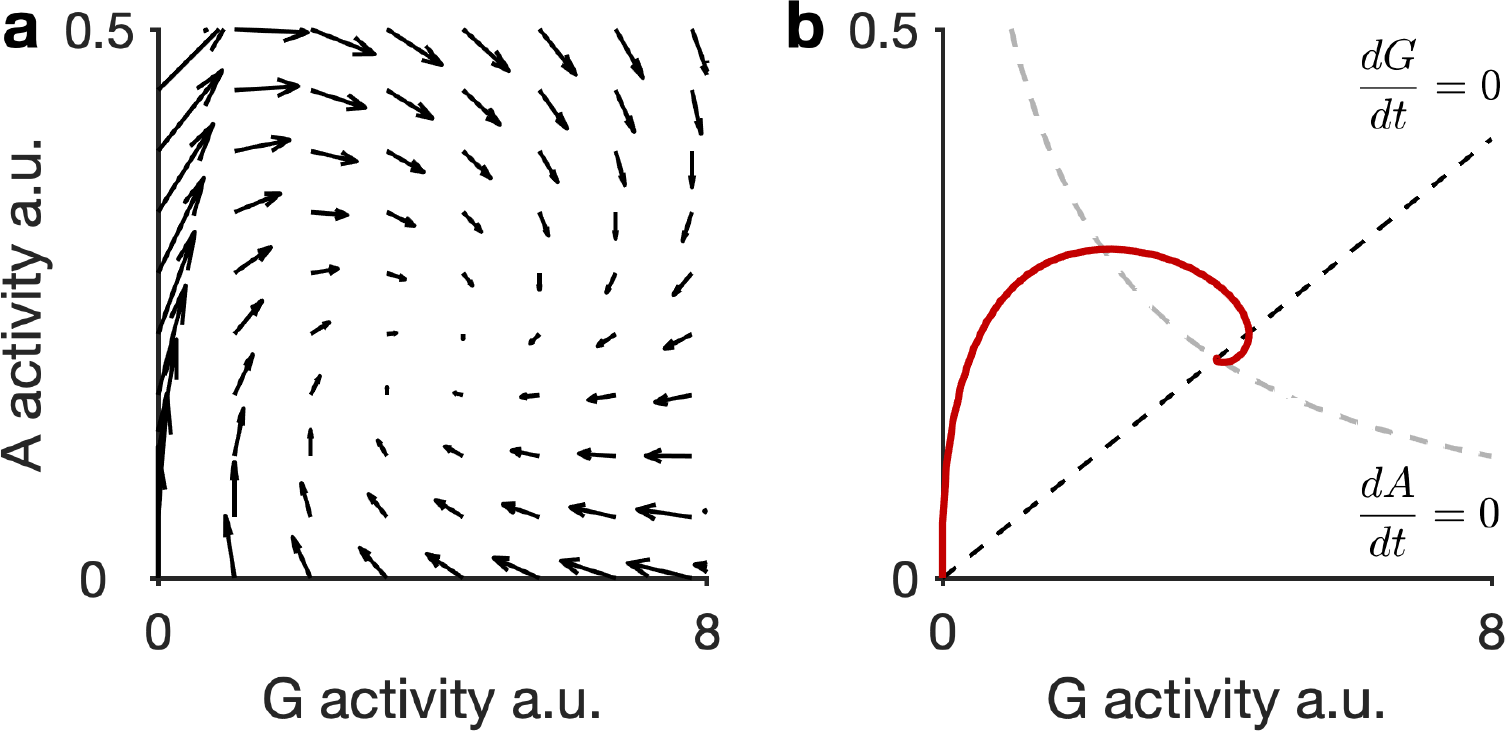
Characteristic model dynamics simulated with *τ_R_* = 1, *τ_G_* = 1, and *ω_I_* = 20. (a) Example model vector field. Arrows indicate instantaneous change in the activities of the (A, G) pair for certain values of *A* and *G*. The vector field shows a stable equilibrium point in the *A*-*G* space. (b) Example network activity trajectory corresponding to the vector field in (a), indicated by red solid line. Grey and black dashed lines indicate the nullclines of the two differential equations for *A* and *G* respectively. The nullclines show values at which either A or G activity does not experience a change in activity (regardless of change in the activity in the other component). The intersection of the two nullclines is where neither the activity of *A* nor *G* changes, again indicating the stable equilibrium point of the network.

